# The *GBA* variant E326K is associated with alpha-synuclein aggregation and lipid droplet accumulation in human cell lines

**DOI:** 10.1101/2022.06.01.494130

**Authors:** Laura. J. Smith, Magdalena. M. Bolsinger, Kai-Yin. Chau, Matthew. E. Gegg, Anthony. H. V. Schapira

**Affiliations:** Department of Clinical and Movement Neurosciences, UCL Queen Square Institute of Neurology, University College London, London, WC1N 3BG; Aligning Science Across Parkinson’s (ASAP) Collaborative Research Network, Chevy Chase, MD, 20815; Division of Medicine, Friedrich-Alexander University Erlangen-Nurnberg, Schloßplatz 4, 91054 Erlangen, Germany

## Abstract

Sequence variants or mutations in the *GBA* gene are numerically the most important risk factor for Parkinson disease (PD). The *GBA* gene encodes for the lysosomal hydrolase enzyme, glucocerebrosidase (GCase). *GBA* mutations often reduce GCase activity and lead to impairment of the autophagy-lysosomal pathway, which is important in the turnover of alpha-synuclein, accumulation of which is a key pathological hallmark of PD. Although the E326K variant is one of the most common *GBA* variants associated with PD, there is limited understanding of its biochemical effects. We have characterised homozygous and heterozygous E326K variants in human fibroblasts. We found that E326K variants did not cause significant loss of GCase protein or activity, endoplasmic reticulum (ER) retention or ER stress, in contrast to the L444P *GBA* mutation. This was confirmed in human dopaminergic SH-SY5Y neuroblastoma cell lines over-expressing GCase with either E326K or L444P protein. Despite no loss of GCase activity, a significant increase of insoluble alpha-synuclein aggregates in E326K and L444P mutants was observed. Notably, SH-SY5Y over-expressing E326K demonstrated a significant increase in lipid droplet number under basal conditions, which was exacerbated following treatment with the fatty acid oleic acid. Similarly, a significant increase in lipid droplet formation following lipid loading was observed in heterozygous and homozygous E326K fibroblasts. In conclusion, the work presented here demonstrates that the E326K mutation behaves differently to common loss of function *GBA* mutations, however lipid dyshomeostasis and alpha-synuclein pathology is still evident.

## Introduction

Mutations or genetic variants in the *GBA* gene (OMIM 606463) are numerically the most important genetic risk factor for Parkinson disease (PD), increasing the prevalence by 5-30-fold depending on the mutation, age and ethnicity (1–5). *GBA* mutation-associated PD (*GBA*-PD) leads to an earlier age of onset and increased cognitive impairment (6–8), with alpha-synuclein pathology similar to that of sporadic PD (6, 9). The *GBA* gene encodes for the lysosomal hydrolase enzyme, glucocerebrosidase (GCase), which catalyses the catabolism of the sphingolipids glucosylceramide (GlcCer) and glucosylsphingosine (GlcSph). Homozygous or bi-allelic *GBA* mutations result in the lysosomal storage disorder Gaucher disease (GD).

Mutations in *GBA* are associated with a specific reduction in GCase activity in the brain (10). The degree of pathogenicity associated with each individual *GBA* mutation differs and variants have been stratified into mild (e.g. N370S) or severe variants (e.g. L444P) (11–13).

*GBA* polymorphic variants like E326K are referred to as risk variants as they do not cause GD when bi-allelic, yet increase the risk for developing PD in both homozygous and heterozygous form (14–16). E326K is one of the most prevalent *GBA* variants in PD patients (17–19) and has been associated with accelerated development of dementia and aggressive motor symptoms (20–22). However, the mechanisms underlying how E326K variants lead to an increased predisposition for PD remain unclear.

The exact mechanisms underpinning the relationship between *GBA* mutations and alpha-synuclein pathology, and how this predisposes some patients to PD, remain elusive. A plethora of models of *GBA*-PD have reported alpha-synuclein pathology (23–28). In cell and animal models of *GBA* mutations activation of the unfolded protein response (UPR) (24, 29–31) and impairment of the autophagy-lysosome pathway (ALP) (23, 24, 32–34) have been demonstrated. In a human neural crest stem-cell derived dopaminergic neuron model of *GBA* mutations, reduced GCase function was accompanied by increased alpha-synuclein levels and could be rescued with the GCase small molecule chaperone ambroxol (28). *GBA* mutations have also been linked to altered lipid metabolism in cell and animal models (23, 27, 35–41), with specific species promoting alpha-synuclein aggregation (42–44). Altered lipid profiles have also been reported in the serum of *GBA*-PD patients (45).

The aim of the present study was to first investigate the effect of the E326K variant on the activity and cellular localisation of GCase in fibroblast cells from patients with homozygous and heterozygous E326K mutations. Fibroblasts homozygous for L444P and N370S were utilised as examples of pathogenic *GBA* mutations of the type that can cause GD. The results revealed that the E326K variant is not associated with a significant loss of GCase activity or protein, unlike L444P and N370S, and does not undergo significant ER-trapping or active the UPR. These findings were confirmed in SH-SY5Y cells expressing E326K, L444P and N370S GCase protein. Despite no loss of GCase activity or activation of the UPR, an accumulation of insoluble alpha-synuclein aggregates was observed in E326K SH-SY5Y cells. A significant elevation in the number of lipid droplets was also demonstrated in fibroblast and SH-SY5Y cells harbouring the E326K variant.

## Results

### No significant reduction in GCase protein levels in E326K mutant fibroblasts

GCase protein level was assessed in fibroblasts by western blot (Figure 1A). In this paper, all results were pooled for genotype. GCase protein and mRNA levels were not significantly different between control and E326K/WT and E326K/E326K lines. In fibroblast lines with homozygous L444P mutations GCase protein was 2.8% of control (p<0.01) and in fibroblasts with homozygous N370S mutations, GCase protein was 17.5% of control (p<0.01). There was no difference in GCase protein levels in young (mean age 9 years) versus older (mean age 61 years) controls (Figure 1B). L444P/L444P fibroblasts exhibited negligible GCase protein levels compared to both.

**Figure 1.**
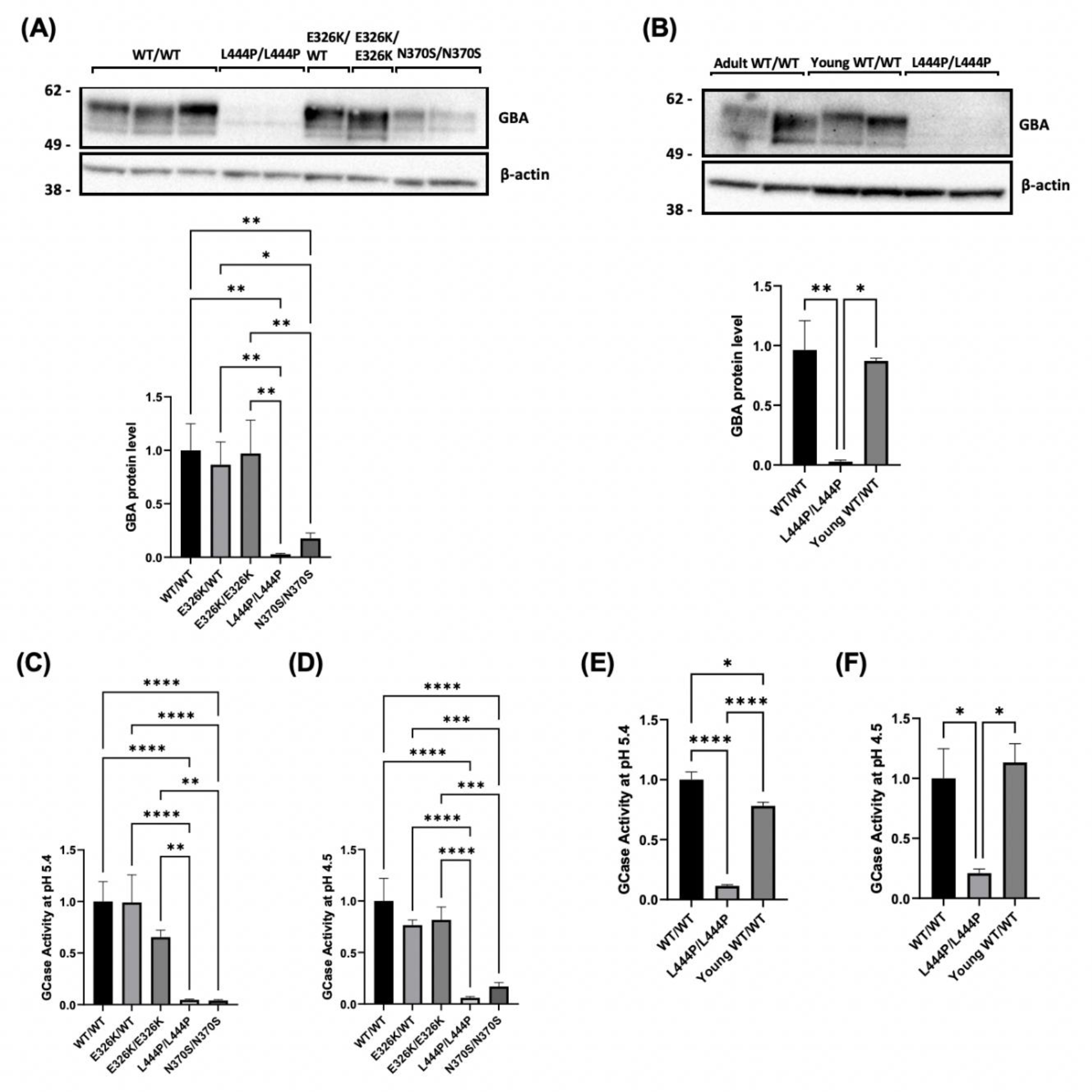
GCase protein level and activity in patient-derived fibroblasts. **(A)** Immunoblot and quantification for GCase protein level in fibroblasts normalised to WT/WT control fibroblasts as shown graphically as mean with SEM (**p<0.01, ***p<0.001, ****p<0.001; n=3). **(B)** Immunoblot and quantification for GCase protein level in adult and young control fibroblasts normalised to WT/WT control fibroblasts (****p<0.0001; n=3). GCase activity assay with M-Glu was performed at **(C)** pH 5.4 with NaT and **(D)** pH 4.5 on fibroblast cell lysates. All data normalised to WT/WT control fibroblasts. GCase activity in in adult and young control fibroblasts at **(E)** pH 5.4 with NaT and **(F)** pH 4.5. (**p<0.01, ***p<0.001, ****p<0.0001; n=4). All data normalised to WT/WT control fibroblasts All graphs show mean with error bars as SEM. Statistical test used was one-away ANOVA with Tukey’s post hoc analysis. Raw data can be found at: 10.5281/zenodo.6553597.

### No significant reduction in GCase enzymatic activity in E326K mutant fibroblasts

Total cell GCase activity was measured at pH 5.4 and revealed no significant changes between controls and E326K/WT (99% of control) and E326K/E326K (65.5% of control) (Figure 1C). A similar pattern was observed at pH 4.5, with E326K/WT cells having 76.6% of control activity and E326K/E326K cells having 82.1% of control activity (Figure. 1D). Both L444P and N370S cells had significantly reduced GCase activity at pH 5.4 (p<0.0001) and pH 4.5 (p<0.0001), compared to controls. Again, results were confirmed in age-matched controls (Figure 1E-1F).

### No significant change in lysosomal content and function in E326K mutant fibroblasts

The overall endo-lysosomal content of fibroblasts was measured by western blot analysis with the lysosomal marker lysosomal-associated membrane protein 1 (LAMP1) (Figure 2A.). No significant alterations were observed between the cell lines. No changes were observed in young and old controls (Figure 2B).

**Figure 2.**
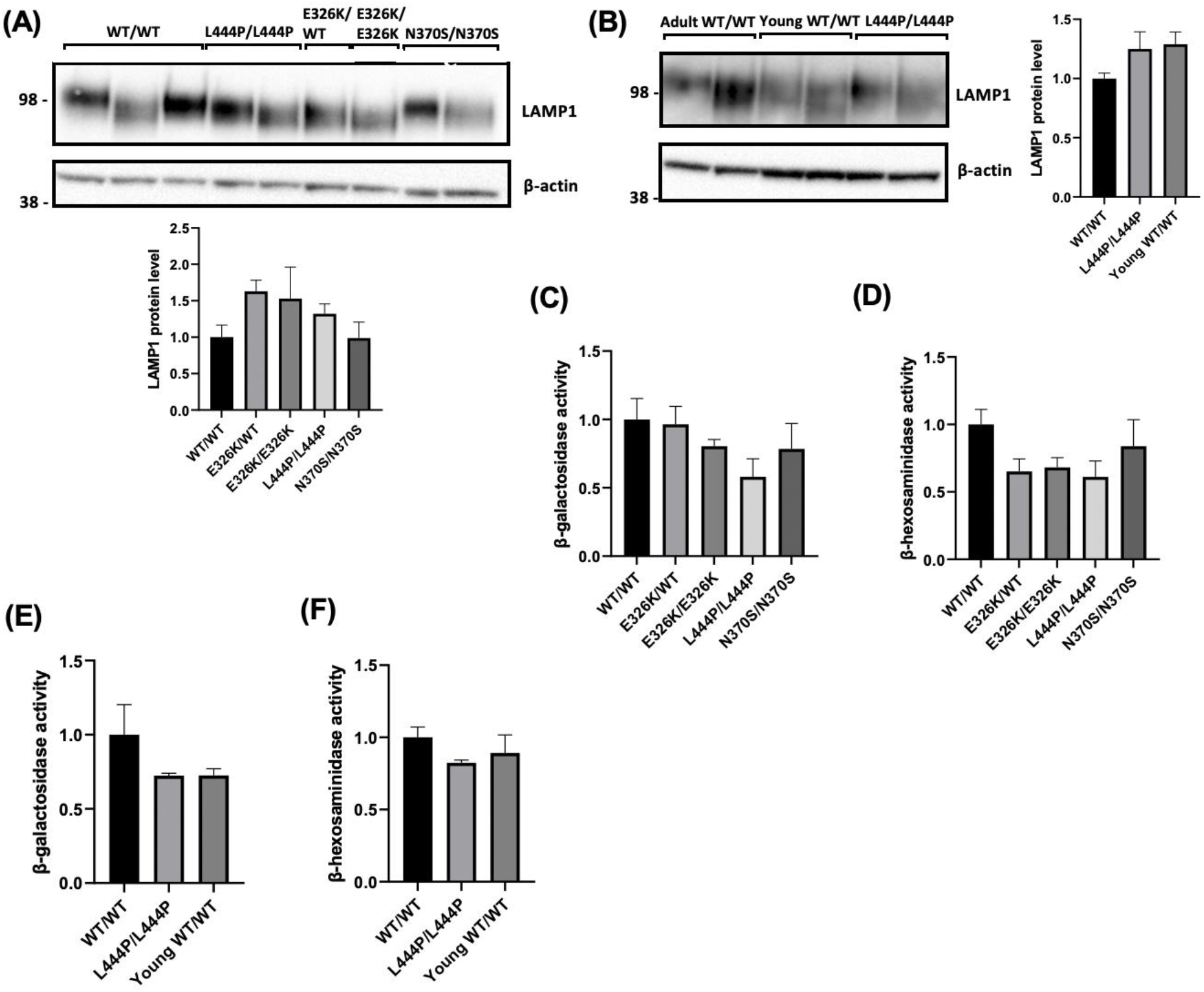
Lysosomal content and function in patient-derived fibroblasts. **(A)** LAMP1 levels were measured via western blot and quantified to assess the overall endo-lysosomal content of the fibroblasts. Data quantified and normalised to WT/WT fibroblast controls. (**B)** LAMP1 levels measured via western blot and compared to young control fibroblasts. Lysosomal function in patient-derived fibroblasts. The activities of lysosomal hydrolases, **(C)** β-galactosidase and **(D)** β-Hexosaminidase were measured at pH 4.1 to assess overall lysosomal function in fibroblasts. All data normalised to WT/WT control fibroblasts. The activities of lysosomal hydrolases, **(E)** β-galactosidase and **(F)** β-Hexosaminidase were measured with young control fibroblasts. All graphs show mean with error bars as SEM. Statistical test used was one-away ANOVA with Tukey’s post hoc analysis. Raw data can be found at: 10.5281/zenodo.6553597.

To investigate overall lysosomal function, enzyme activity assays were performed to measure the activity of two other lysosomal hydrolases, β-galactosidase and β-hexosaminidase (Figure 2C-3F). No significant changes were observed between control lines and mutant fibroblasts. Analysis in age-matched controls revealed no significant alterations between aged and young controls.

**Figure 3.**
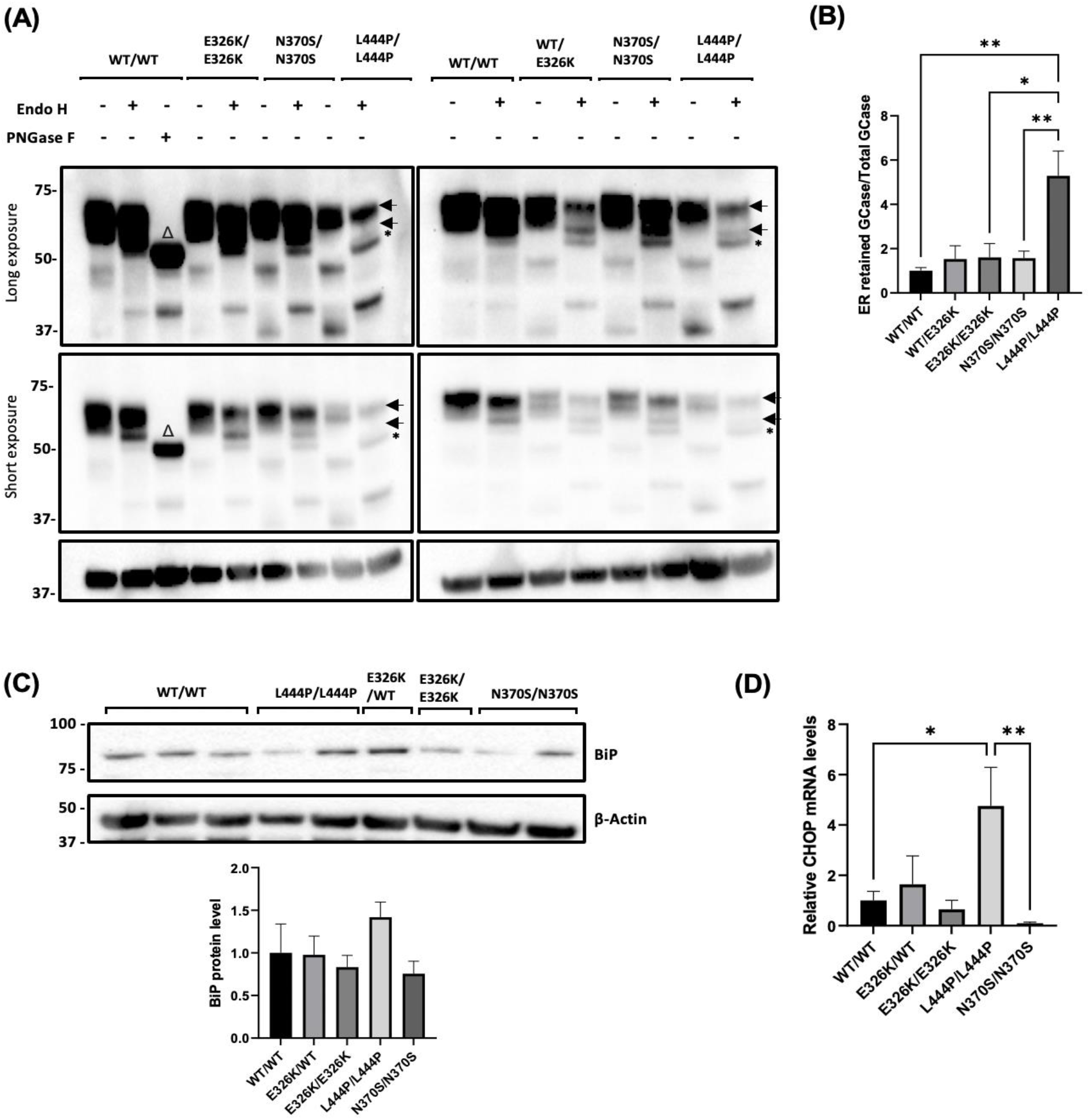
ER retention and ER stress in patient-derived fibroblasts. **(A)** Fibroblast cell lysates (20 μg of WT, E326K and N370S mutants and 70 μg of L444P mutants) were treated with or without endoglycosidase-H (Endo H) and GCase protein species analysed by Western blot. WT/WT cell lysate (20 μg) was treated with Peptide-N-Glycosidase F (PNGase F) as a positive control. Figure shows blots at long and short exposure. The two normal species of GCase detected in fibroblasts are indicated by arrows. An additional lower molecular weight band was observed, indicating ER retained GCase, in L444P/L444P fibroblasts following endo-H treatment (black asterisk). The WT/WT cell line treated with PNGase exhibited a lower molecular weight band, indicating a GCase species with no glycosylation (Δ). **(B)** Quantitative analysis of Endo H digestion displayed as the density of bands for ER resident GCase divided by the density of bands for total GCase protein, normalised to β-actin band density. Data normalised to WT/WT which was set at 1. ER stress was analysed by **(C)** quantifying BiP protein level in fibroblasts. Results are normalized to WT/WT control fibroblasts. **(D)** The expression of *CHOP* mRNA levels in patient fibroblasts was quantified and normalized to WT/WT controls. For each experiment, two biological replicates were used for each cell line. For quantification, *CHOP* expression for each cell line was calculated, pooled and averaged for each genotype. Graph shows mean and error bars show SEM. Statistical test used was one way ANOVA with Tukey post-hoc analysis (*p<0.05, **p<0.01; n=3). Raw data can be found at: 10.5281/zenodo.6553597.

### No significant ER retention of GCase in E326K mutant fibroblasts

The different glycosylation patterns of GCase, as it passes through the secretory pathway, can be utilised to assess the proportion of GCase that is trapped in the ER. ER-retained forms of GCase carry N-linked glycans that are sensitive to cleavage by Endo-H, producing lower molecular bands on western blot. The GCase protein resolves as 3 bands after treatment, with the lower exposure blot showing two higher molecular weight bands likely to be mature protein (Figure 3A). For analysis, wild-type fibroblasts were digested by PNGase F as a positive control for unglycosylated GCase (Δ symbol on blot). The corresponding ER-resident bands in the mutant cell lines are marked with a black asterisk. This band is absent in wild-type cell lines. To quantify the retention of mutant GCase protein, the density of the ER-resident band was divided by the density of the two mature GCase bands (Figure 3B). Compared to controls, L444P/L444P fibroblasts were the only cells to exhibit an increase in Endo-H sensitive GCase fraction (5.28-fold) (p<0.01).

To further investigate GCase trafficking, lysosomal GCase activity was measured using a substate that is taken up only in to acidic vesicles and fluoresces upon catalysis with GCase (46). Lysosomal GCase activity was measured in E326K/+, E326K/E326K and L444P/L444P fibroblasts (Supplemental Figure 1). After loading of substrate, there is a linear increase in fluorescent product up to 45 minutes, after which GCase enzymatic activity plateaus. This plateau is the depletion/release of substrate from late endosomes and lysosomes over time. Therefore, the initial linear rate of enzyme activity was calculated between time 0 and time 45 minutes. Lysosomal GCase activity was abolished in wild-type fibroblast cells pre-treated with 10 μM CBE for 24 hours, a GCase specific inhibitor, and was similar to lysosomal GCase activity in L444P/L444P fibroblasts. In comparison, E326K/WT and E326K/E326K fibroblasts have activity in the lysosome closer to that of controls, suggesting this variant behaves differently to the L444P loss of function variant.

### No significant ER stress in E326K mutant fibroblasts

To assess UPR activation in fibroblasts, BiP chaperone protein levels were measured (Figure 3C). BiP (GRP78) is a central regulator of the UPR, binding to unfolded proteins in the ER and activating the three arms of the UPR. Expression levels can also be increased during ER stress (47). No significant changes were observed in any mutant lines compared to healthy controls.

Analysis of *CHOP* mRNA expression, which is up-regulated in response to prolonged activation of the UPR following binding of BiP to the PERK arm of the UPR (48) revealed a 4.75-fold increase in L444P/L444P fibroblasts (p<0.05) (Figure 3D). No significant increases were observed in E326K or N370S fibroblasts compared to controls.

### Characterisation of GCase protein level, expression and activity in SH-SY5Y cell lines expressing GCase mutants

SH-SY5Y stable cell lines expressing wild-type, E326K, L444P and N370S *GBA* constructs were generated and data from two clones pooled for each genotype. Over-expression of *GBA* was confirmed via qPCR (Figure 4A). Wild-type (288-fold), E326K (190-fold), L444P (209-fold) and N370S (125-fold) lines all expressed similar higher levels of GCase mRNA, when compared to untransfected cells.

**Figure 4.**
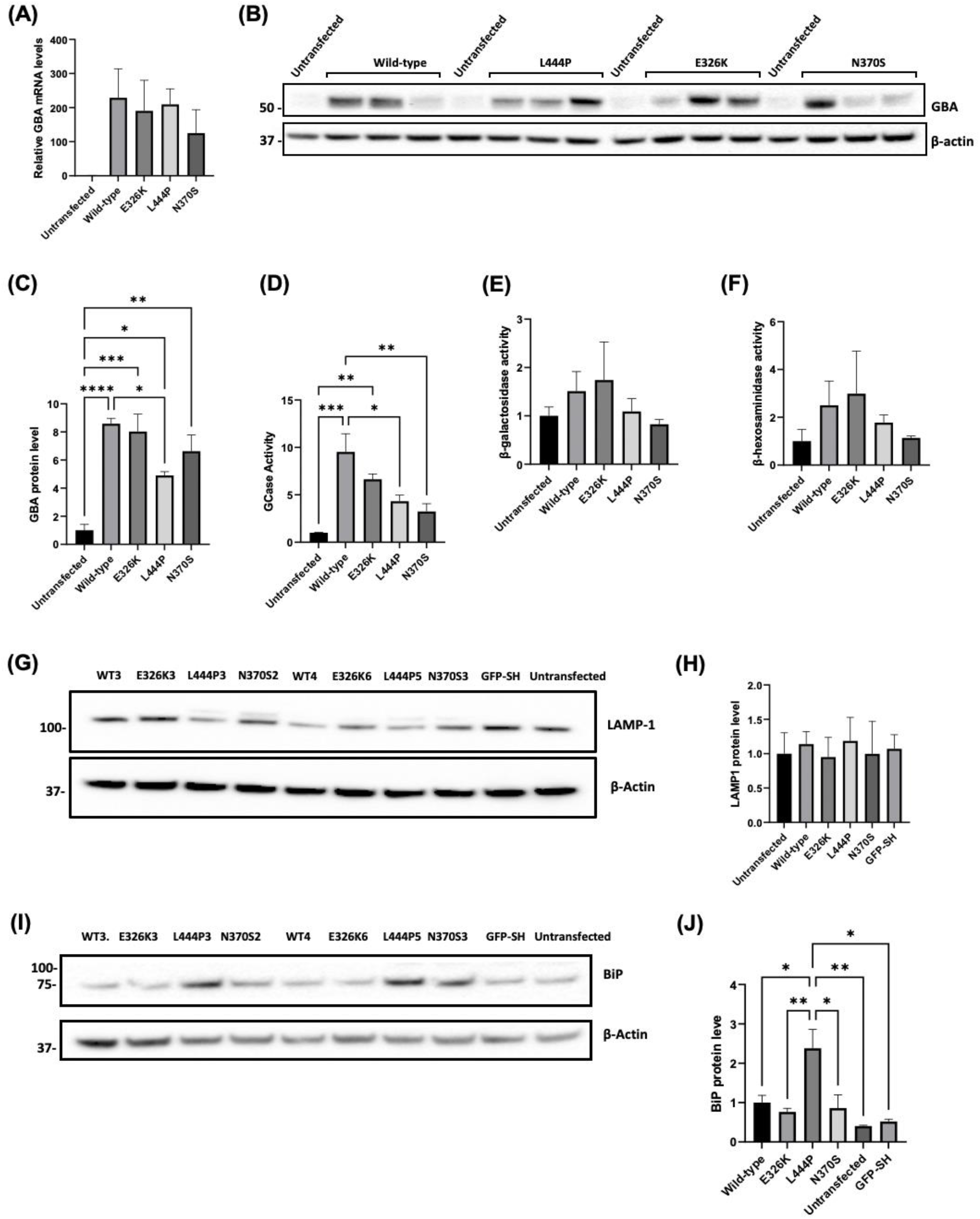
*GBA* levels, lysosomal function and ER stress in SH-SY5Y cells over-expressing mutant GCase. **(A)** Quantification of *GBA* mRNA levels in stable SH-SY5Y cell lines, normalised to untransfected SH-SY5Y cells (set at 1). **(B)** Immunoblot and **(C)** quantification of *GBA* protein levels in stable SH-SY5Y cell lines. Data normalised to untransfected SH-SY5Y cells (set at 1). Graphs show mean with SEM. Statistical test used was one-way ANOVA with Tukey’s post hoc analysis (*p<0.05, **p<0.01, ***p<0.001, ****p<0.0001; n=4). **(D)** GCase activity in nmole/hr/mg in stable SH-SY5Y cell lines measured at pH 5.4 with NaT and normalised to untransfected SH-SY5Y cells (set at 1) Graphs show mean with SEM. Statistical test used was one-way ANOVA with Tukey’s post hoc analysis (*p<0.05, **p<0.01, ***p<0.001; n=4). The activities of lysosomal hydrolases, β-galactosidase and β-Hexosaminidase were measured at pH 5.1 in SH-SY5Y stable cell lines. **(E)** β-galactosidase activity in nmole/0.5hr/mg in undifferentiated SH-SY5Y clones normalised to untransfected SH-SY5Y cells. **(F)** β-Hexosaminidase activity in nmole/0.5hr/mg measured in undifferentiated SH-SY5Y clones normalised to untransfected SH-SY5Y cells. **(G)** LAMP1 levels were measured via western blot and **(H)** quantified to assess the overall endo-lysosomal content of the SH-SY5Y cell lines. Immunoblot and quantification for LAMP1 protein level in SH-SY5Y stable cell lines. Data normalised to wild-type SH-SY5Y cells (n=3). **(I)** Immunoblot and **J)** quantification of BiP protein level in SH-SY5Y clones. Data normalised to untransfected SH-SY5Y cells. Graphs show mean with SEM. Statistical test used was one-way ANOVA with Tukey’s post hoc analysis(*p<0.05. **p<0.01; n=3). Raw data can be found at: 10.5281/zenodo.6553597.

Compared to untransfected SH-SY5Y cells, all GCase over-expressing lines had higher GCase protein levels and activity (p<0.05-p<0.0001) (Figure 4B-4D). No significant changes in GCase protein level or activity were observed between wild-type and E326K cells. Cells expressing L444P mutant protein had lower GCase protein levels (55.4% of wild-type) (p<0.05) and activity (45.5% of wild-type) (p<0.05) compared to wild-type. In N370S cells, GCase protein level was not significantly reduced unlike activity, which, but activity was 33.6% of wild-type (p<0.01).

### Overall lysosomal content and function unaltered by GCase mutations in undifferentiated SH-SY5Y cells

No significant changes were observed in activity of the lysosomal hydrolases, β-galactosidase and β-Hexosaminidase, and protein level of LAMP1 across SH-SY5Y cell lines (Figure 4E-4H). These data indicate that *GBA* mutations do not influence the overall endolysosomal content in SH-SY5Y cells.

### Increased ER stress in undifferentiated SH-SY5Y cells over-expressing L444P mutant GCase

Activation of the UPR in SH-SY5Y cells was investigated by measuring BiP protein levels. Compared to wild-type, cells expressing L444P GCase protein were the only *GBA* lines that exhibited a significant elevation in BiP levels (p<0.05) (Figure. 4I-4J). This was also significantly higher than E326K (p<0.01) and N370S (p<0.05) lines. As a control, cells over-expressing green fluorescent protein, which has a similar size to GCase, was also not found to increase BiP levels. Note that *CHOP* mRNA levels in SH-SY5Y were extremely low (> 30 cycles) and could not be measured reliably.

### Increased soluble intracellular alpha-synuclein protein level in undifferentiated SH-SY5Y cells over-expressing L444P mutant GCase

Accumulation of alpha-synuclein monomers in cells is a key feature in models of *GBA*-PD (23, 26, 33, 34, 39, 46, 49). Analysis of Triton X-100 soluble, monomeric alpha-synuclein revealed no significant changes in E326K and N370S cells compared to wild-type (Figure 5A-5B), while those expressing the L444P variant exhibited a 2.58-fold increase (p<0.05). There were no significant changes in *SNCA* mRNA expression observed between cell lines (Figure 5C).

**Figure 5.**
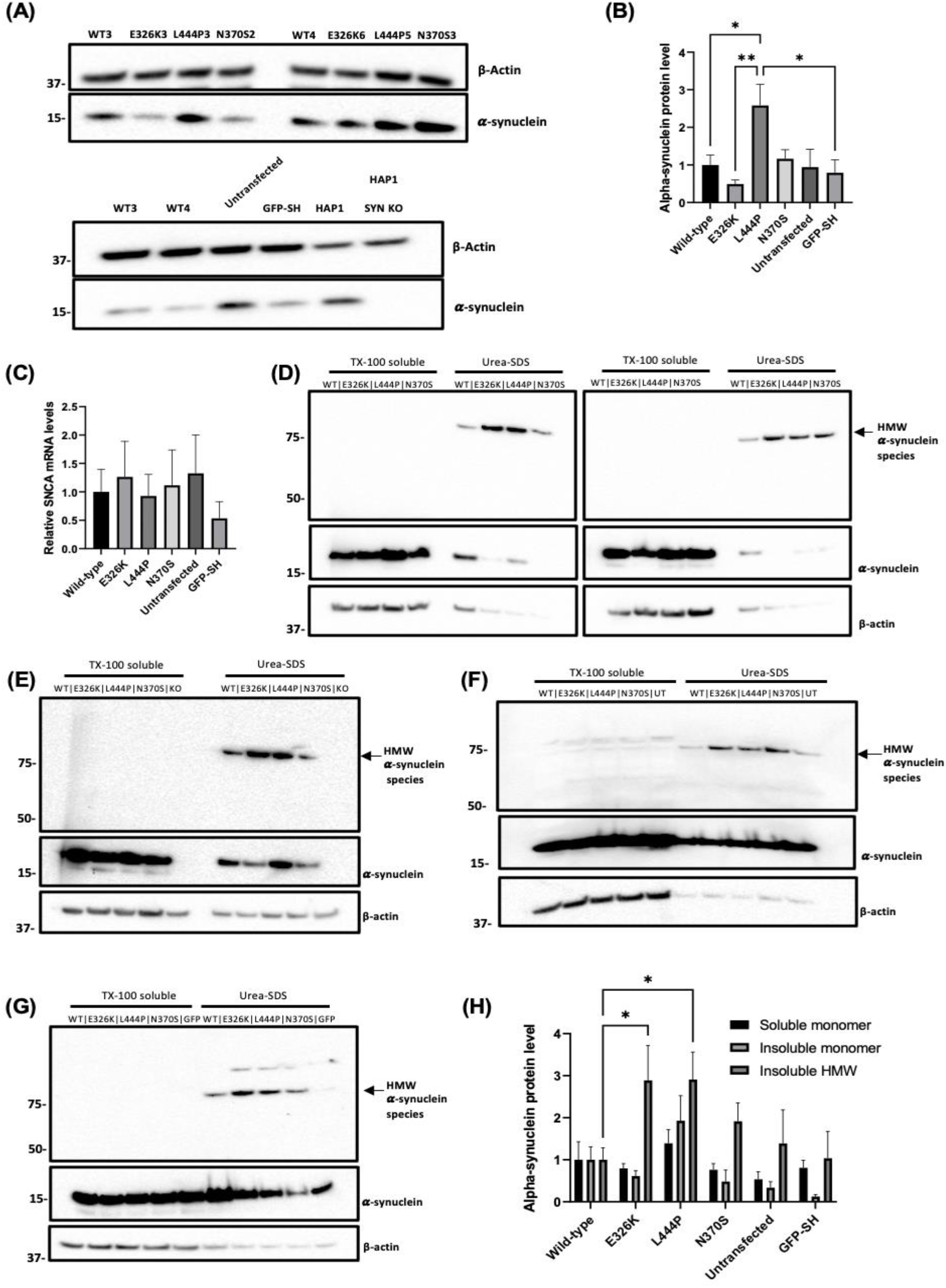
Soluble and insoluble alpha-synuclein levels in SH-SY5Y stable cell lines expressing mutant *GBA*. **(A)** Immunoblot of alpha-synuclein protein level in SH-SY5Y clones. **(B)** Quantification of alpha-synuclein immunoblotting in SH-SY5Y clones normalised to wild-type clones. **(C)** Quantification of *SNCA* mRNA levels in SH-SY5Y stable clones normalised to wild-type clones (n=3). TX-100 soluble and insoluble fractions (urea-SDS) were made from cells and analysed for alpha-synuclein by western blotting. HMW alpha-synuclein species (arrow) were detected in urea-SDS fractions. Monomeric alpha-synuclein was present in the TX-100 soluble fraction and some urea-SDS fractions. **(D)** Immunoblot for all SH-SY5Y cell lines over-expressing mutant *GBA* protein. Appropriate control cell lines were also analysed: **(E)** HAP1 alpha-synuclein knock-out cells (KO); **(F)** untransfected SH-SY5Y cells (UT) and **(G)** GFP over-expressing SH-SY5Y cells (GFP). **(H)** Quantification of soluble and insoluble alpha-synuclein immunoblots. Data normalised to wild-type clones. Ten technical repeats. Graphs show mean with SEM. Statistical test used was one-way ANOVA with Tukey’s post hoc analysis or two-way ANOVA with Tukey’s post hoc analysis (*p<0.05; n=16). Raw data can be found at: 10.5281/zenodo.6553597.

### Increased insoluble alpha-synuclein protein level in undifferentiated SH-SY5Y cells over-expressing mutant GCase protein

As alpha-synuclein can form insoluble, phosphorylated aggregates (50), the effect of *GBA* mutations on alpha-synuclein aggregates in the Triton X-100 insoluble fraction was investigated. Soluble and insoluble Triton X-100 fractions were run on the same western blot and immunoblotted for alpha-synuclein (Figure. 5D-5H). The vast majority of alpha-synuclein remained in the soluble fraction. Monomeric, soluble alpha-synuclein was a single band at the expected molecular weight of 15 kDa. No significant changes in soluble alpha-synuclein monomers were observed.

There was evidence of a HMW alpha-synuclein species in the insoluble fraction (75 kDa), which was absent in human *SNCA* knock-out HAP1 lines (Figure. 5E). Insoluble alpha-synuclein band density was expressed as a ratio against soluble β-actin (Figure. 5H). As expected in the insoluble fraction, β-actin band density was lower than the soluble fraction and likely represents β-actin associated with Triton X insoluble membranes (e.g., lipid rafts) There was a concurrent and significant increase in E326K (2.88-fold) and L444P (2.91-fold) cells compared to wild-type (p<0.05), indicative of an accumulation of insoluble alpha-synuclein aggregates.

### Increased number of lipid droplets in E326K mutant cell lines

As *GBA* mutations have been associated with altered lipid metabolism (51–53), lipid droplet accumulation was measured in our cell lines. Lipid droplets are organelles involved in intracellular lipid homeostasis (54). BODIPY 493/503 stains neutral lipids and lipid droplets are visualised as punctate structures, examples of which are shown by arrows (Figure. 6). The number of lipid droplets were quantified using Image J and normalised to cell area. To further analyse the effect of *GBA* mutations on lipid droplet accumulation, SH-SY5Y cells were grown under basal conditions and stained with BODIPY 493/503 (Figure. 6D-6E). In these conditions, a higher number of lipid droplets was observed in mutants compared to wild-type, significantly elevated in E326K (2.1-fold) (p<0.0001; n=3) and L444P (1.8-fold) (p<0.0001). Both were significantly higher compared to N370S expressing lines (E326K p<0.0001 and L444P p<0.01). The number of lipid droplets in untransfected cells was significantly lower than all over-expressing cell lines (wildtype p<0.05; E326K, L444P and N370S p<0.0001). However, SH-SY5Y cells over-expressing GFP were not significantly different to untransfected cells, suggesting this increase in lipid droplet number is not simply an artefact of protein over-expression.

**Figure 6.**
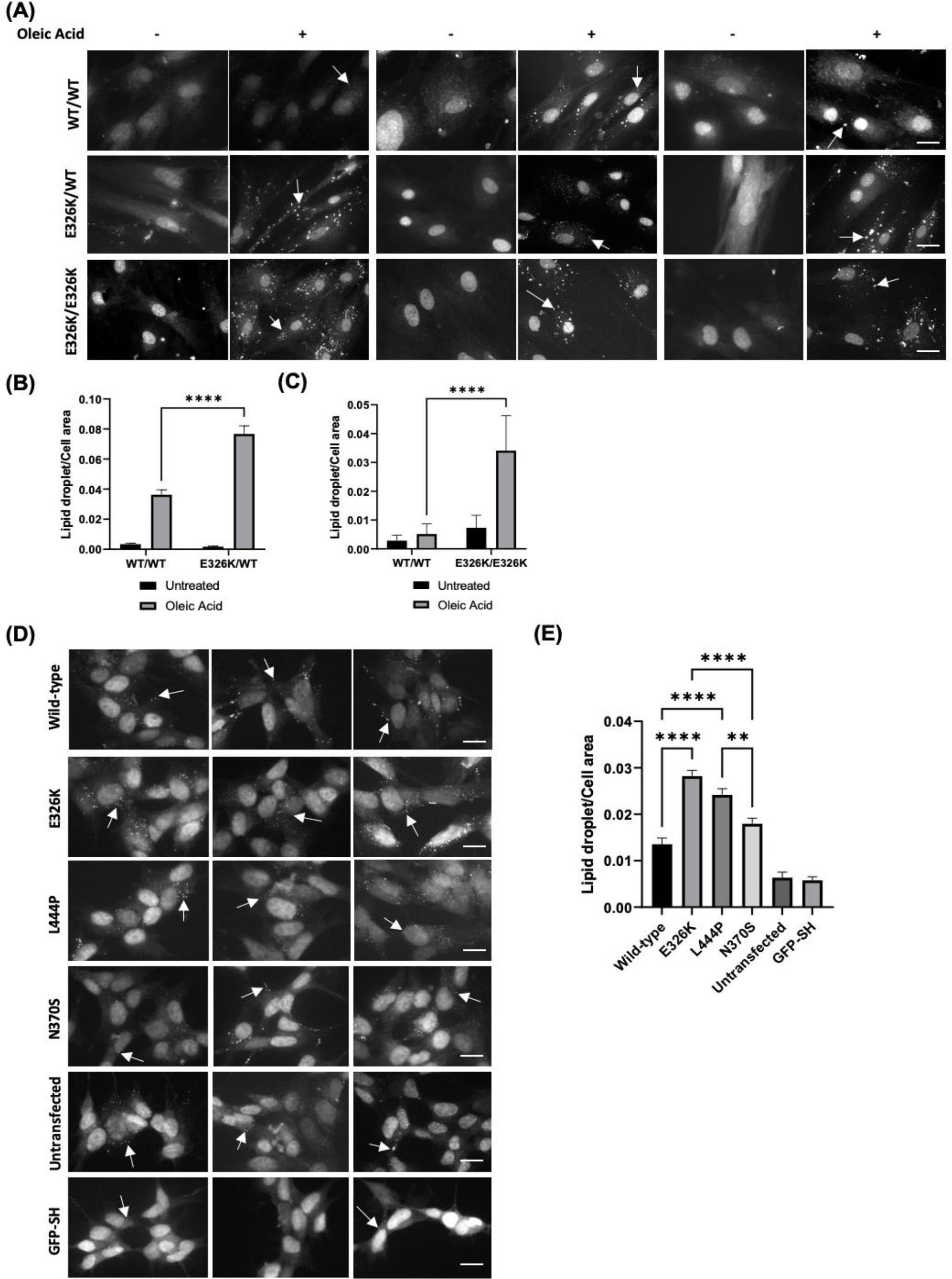
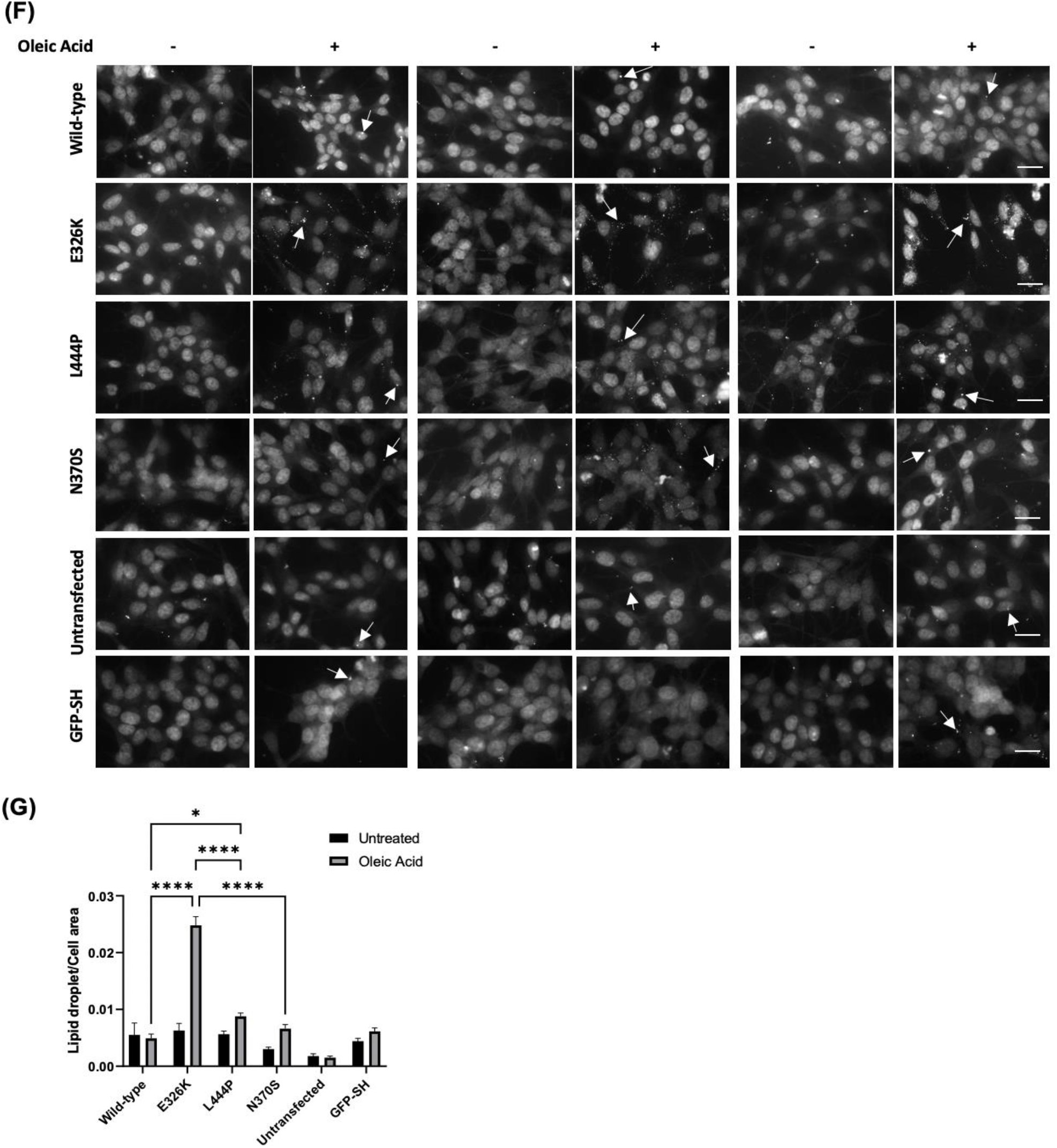
Lipid droplets in *GBA* mutant cells. **(A)** Fibroblast lines harbouring the E326K mutations were starved in Opti-MEM overnight and incubated with and without 100 µM oleic acid for 5 hours. Cells were stained with lipophilic fluorescent probe BODIPY 493/503. Lipid droplets present as punctate structures. Three representative images from each genotype shown (-) denotes untreated and (+) denotes treated with oleic acid. All cells were counted in the images and the number of lipid droplets counted and normalised to cell area using ImageJ. Scale bar represents 20 µm. Quantification of lipid droplets in **(B)** control and E326K/+ fibroblasts and **(C)** control and E326K/E326K fibroblasts displayed as mean with error bars showing SEM. Statistical test used was one-way ANOVA with Tukey post-hoc analysis or two-way ANOVA with Tukey’s post-hoc analysis (****p<0.0001). **(D)** SH-SY5Y cells over-expressing mutant GCase protein were grown on coverslips and stained with lipophilic fluorescent probe BODIPY 493/503. Lipid droplets present as punctate structures. Three representative images from each genotype shown. All cells were counted in the images and the number of lipid droplets counted and normalised to cell area using ImageJ. **(E)** Quantification of lipid droplets per cell area shown as mean with SEM. Statistical test used was one-way ANOVA with Tukey post-hoc analysis or two-way ANOVA with Tukey’s post-hoc analysis (**p<0.01; ****p<0.0001). **(F)** SH-SY5Y cells over-expressing mutant GCase protein were starved in Opti-MEM overnight and incubated with and without 100 µM oleic acid for 5 hours. Cells were stained with lipophilic fluorescent probe BODIPY 493/503. Lipid droplets present as punctate structures. Three representative images from each genotype shown (-) denotes untreated and (+) denotes treated with oleic acid. All cells were counted in the images and the number of lipid droplets counted and normalised to cell area using ImageJ. **(G)** Quantification of lipid droplets per cell area displayed as mean with error bars showing SEM. Statistical test used was one-way ANOVA with Tukey post-hoc analysis or two-way ANOVA with Tukey’s post-hoc analysis (*p<0.05; ****p<0.0001). Scale bar represents 20 µm. Raw data can be found at: 10.5281/zenodo.6553597.

To investigate if the increased LD number in *GBA* mutant cells was due to increased rate of lipid droplet formation, SH-SY5Y cells were starved overnight in OptiMEM and then treated with the fatty acid oleic acid for 5 hours to induce lipid droplet formation (55) (Figure 6F-6G). A 5.1-fold increase in lipid droplet formation was observed in E326K cells compared to wild-type cells (p<0.0001). In SH-SY5Y cell lines expressing L444P GCase protein a 1.77-fold increase in lipid droplets, compared to wild-type, was observed (p<0.05). Lipid droplet formation in E326K cells was significantly higher than in L444P (p<0.0001), N370S (p<0.0001) and GFP (p<0.0001) lines. Both E326K and L444P were significantly higher than untransfected (E326K p<0.0001 and L444P p<0.001).

To confirm the role of E326K in LD formation, fibroblast lines heterozygous and homozygous for E326K were starved overnight, loaded with oleic acid and lipid droplet formation quantified (Figure 6A-6C). Compared to control cells, E326K/WT cells exhibited a 2.11-fold increase in lipid droplet formation following lipid loading with oleic acid (p<0.0001) (Figure 6B). Similarly, following oleic acid treatment, a 6.61-fold increase in lipid droplet formation was observed in E326K/E326K cells compared to controls (Figure 6C).

## Discussion

The results of this study indicate that the E326K variant does not behave in a similar fashion to the pathogenic *GBA* mutations L444P and N370S. Notably, despite no significant loss of E326K GCase activity, insoluble alpha-synuclein aggregates in SH-SY5Y cells were observed and were coincident with an enhanced formation and accumulation of lipid droplets in SH-SY5Y and fibroblast lines, suggestive of altered lipid homeostasis. A schema highlighting the potential pathways underlying the mechanism of the E326K variant is in Figure 7.

**Figure 7.**
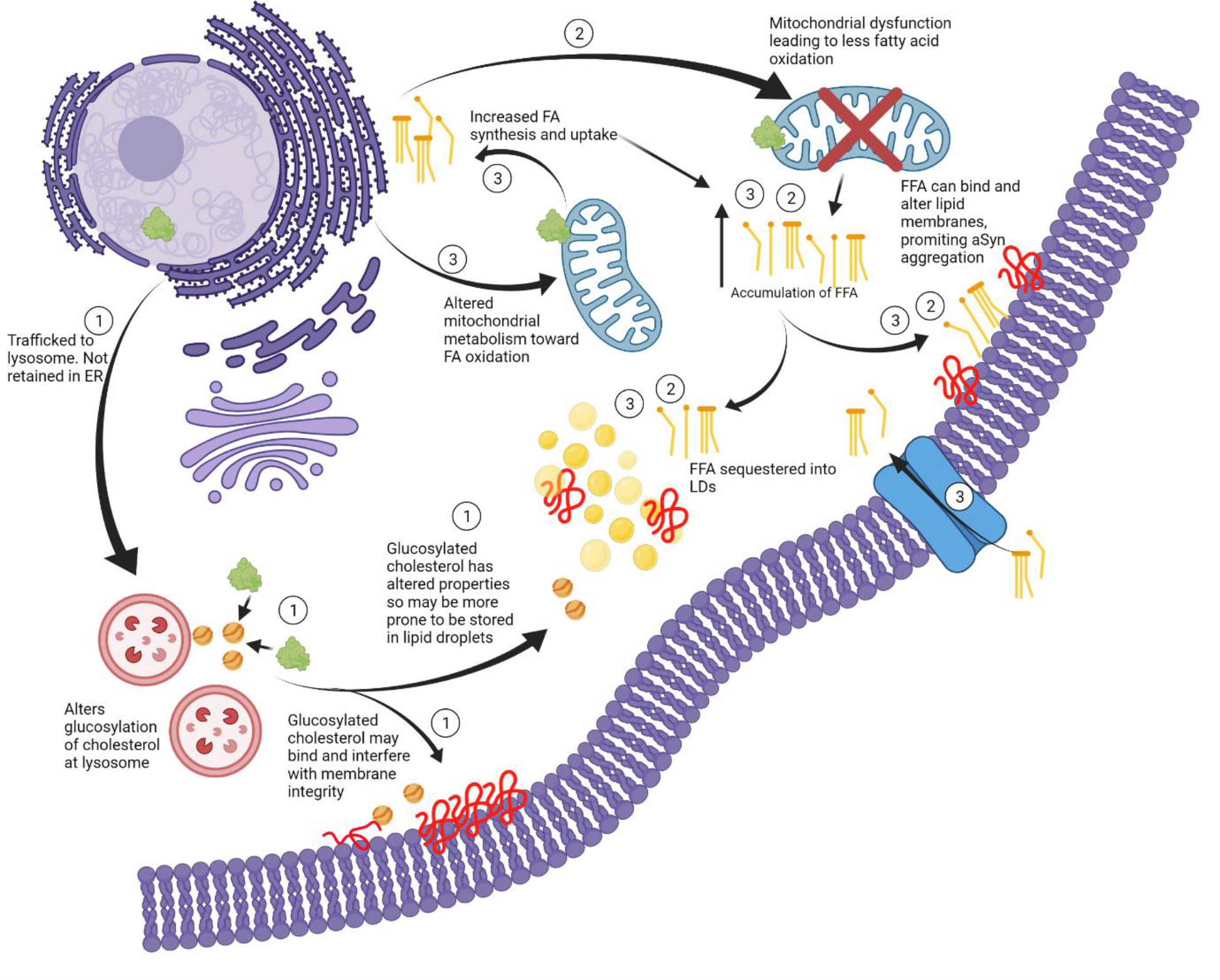
Proposed mechanisms underlying E326K GCase pathology. **(A)** E326K GCase is likely not retained in the ER and is trafficked to the lysosome. At the lysosome GCase carries out its ‘moonlighting’ function of glucosylating cholesterol. E326K GCase may have increased or decreased ability to glucosylate cholesterol, which alters the properties of accumulated cholesterol. This cholesterol can interfere with lipid membranes, altering lipid composition, and promoting alpha-synuclein aggregation. This cholesterol has altered properties so may be more prone to storage in to lipid droplets, leading to an accumulation of lipid droplets. Lipid droplets can act as a site to bind alpha-synuclein, and in high concentrations and under pathological conditions alpha-synuclein may aggregate at lipid droplets. **(2)** The E326K GCase mutation may lead to dysfunctional mitochondria, potentially through impaired clearance. These dysfunctional mitochondria may have reduced ability to metabolise fatty acids, leading to the accumulation of free fatty acids in the neuron. Free fatty acids are capable of interfering with lipid membranes, potentially promoting alpha-synuclein aggregation. Free fatty acids are also sequestered into lipid droplets, as the neuron attempts to protect the cell from lipotoxicity. **(3)** The E326K mutation may cause a shift in the mitochondria’s metabolism capacity, shifting away from glycolysis toward fatty acid oxidation to provide energy for the neuron. In order to meet the demand for fatty acids, the neuron may be more primed to synthesise fatty acids or take up external fatty acids. This may lead to an accumulation of free fatty acids in the neuron, which can exert lipotoxic effects, accumulate in lipid droplets and may induce alpha-synuclein aggregation.

Few studies have biochemically characterised E326K GCase protein. This study provides evidence that E326K in fibroblasts does not induce a significant loss of GCase function or result in unfolded protein response, unlike other common pathogenic *GBA*-PD mutations N370S and L444P. In homozygous and heterozygous form, E326K mutations are not associated with a significant loss of GCase expression or activity in fibroblasts and SH-SY5Y cells. These findings are supported by previous literature demonstrating the E326K variant generally reduces GCase activity to a lesser extent than other pathogenic *GBA* mutations (16, 56–61), while L444P and N370S variants are reported to induce a loss of GCase function (30, 31, 42, 61–67), as demonstrated in this study.

Aberrant trafficking and retention of mutant GCase in the ER has been reported in a variety of cell models (30, 31, 61, 62, 68–71). Activation of the UPR has been demonstrated extensively in cells harbouring the L444P variant (29, 31, 61, 68, 70, 71). In this study, unlike L444P, the E326K protein is not localised to the ER in fibroblasts and does not activate the UPR, suggesting correct lysosomal trafficking and the absence of a severely misfolded protein. It may be that the mutant E326K protein exerts a gain of function mechanism through a ‘moonlighting’ function such as the glucosylation of cholesterol at the lysosome (72). The altered properties of glucosylated cholesterol may increase its storage in lipid droplets or influence lipid membrane fluidity, which may promote the aggregation of alpha-synuclein (73). Alterations to the composition of lipid rafts, which are central regulators of CMA (74), may impair CMA-mediated degradation of alpha-synuclein. Furthermore, an increase in cholesterol has been associated with an impairment of autophagy in N370S fibroblasts (67). As ER retention of GCase is reported to correlate with disease severity (75) and has been shown to be variable among lines harbouring the same N370S genotype (70), this may explain the lack of impaired GCase localisation in N370S cells in this study.

Evidence suggests a role for *GBA* mutants in the accumulation of alpha-synuclein monomers and aggregates (76). Despite no loss of GCase activity, SH-SY5Y cells expressing the E326K variant exhibit an increase in insoluble alpha-synuclein aggregates. This was increased to a similar level as L444P mutant cells. As the L444P variant has been shown to accelerate alpha-synuclein pathology and spread in mice (77, 78), this points toward a potentially similar propagation of pathology in E326K carriers. The observation that the E326K mutation occurs on the GCase protein surface (79), suggests protein interactions may be influenced. It’s possible that the E326K mutant GCase protein has a reduced affinity for binding alpha-synuclein, as suggested in the N370S protein (80), which could induce alpha-synuclein lipid binding and aggregation.

Accumulation of insoluble alpha-synuclein aggregates may be explained by the accumulation of lipid droplets in E326K fibroblasts and SH-SY5Y cells, suggestive of altered lipid metabolism. Lipid homeostasis is necessary for maintaining proper function of the neuron and synaptic plasticity. Alterations in such pathways have been reported in PD with increased levels of TAGs, cholesterol and ceramides (81). In an effort to overcome lipid overload, cells initially induce a pathway to store excess lipids in lipid droplets (54). Although suggested to be initially protective (82, 83), excess lipid droplets can be neurotoxic (84). Lipid droplets protect cells from excess fatty acids which can be targeted for lipid peroxidation. During oxidative stress, reactive oxygen species (ROS) can attack free fatty acids and generate toxic lipid peroxides and reactive aldehydes which can exert lipotoxicity including damage to lipid membranes, ER stress, mitochondrial damage and subsequent neurodegeneration (82, 83, 85–88). Excessive lipid droplet formation has however been demonstrated to trigger neurotoxicity (84). It may be this that contributes to neurodegeneration in E326K mutants.

Under normal conditions, the number of lipid droplets accumulating in SH-SY5Y cells expressing E326K and L444P was elevated compared to wild-type cells. Following lipid loading, lipid droplet formation was significantly elevated in these cells compared to wild-type and other mutant lines. The same was observed in heterozygous and homozygous E326K fibroblasts. Interestingly, although L444P SH-SY5Y cell lines exhibited increase lipid droplets after lipid loading, this was to a much lesser extent than E326K lines. As the E326K variant is not associated with a loss of GCase function or activation of the UPR, yet the L444P line is, it is likely that these variants induce accumulation of lipid droplets through different pathways. The L444P variant may lead to improper ALP and clearance of lipids, whereas the E326K variant may work through pathways involving the mitochondria and metabolism of lipids.

Many studies demonstrate a correlation between increased lipid droplets and alpha-synuclein pathology (83, 89, 90). Recently, in a mouse model of synucleinopathy, lipid droplet accumulation correlated with alpha-synuclein pathology, both of which were rescued by over-expressing wild-type GCase (91), reinforcing the role of GCase in this pathway. Proper regulation of cellular lipids is critical to maintain the composition and fluidity of lipid membranes. Such lipids can bind alpha-synuclein and accelerate its formation into toxic oligomeric and fibrillar species, propagating PD pathology. In addition, alterations in lipid membrane integrity can influence alpha-synuclein binding and enhance the conversation of monomeric alpha-synuclein into toxic aggregates (81). Altered composition of GSLs has been demonstrated in L444P and N370S neurons levels (23, 24, 44), although the accumulation of these lipids in *GBA*-PD is debatable (92, 93). A shift toward short-chain GSLs may be induced by *GBA* mutations which can promote the toxic aggregation of alpha-synuclein (42).

The E326K mutation may influence lipid metabolism, possibly inducing its remodelling toward fatty acid oxidation, priming neurons to more efficiently synthesise or take up fatty acids. Fatty acid oxidation occurs in the mitochondria (94) and a defective mitochondrial network may contribute to E326K pathogenesis (95). Mitochondrial dysfunction has long been implicated in the pathogenesis of PD and associated with *GBA* mutations (32, 96–98). There may also be a reciprocal relationship between mitochondria and lipid droplets as they may associate and form mitochondria with a unique structure and function less prone to fatty acid oxidation (95).

Understanding the effect of the E326K variant in human midbrain dopamine neurons is an exciting avenue to explore in future work, including the potential role of mitochondria in the mechanism of disease. This may shed light on why the E326K variant contributes to the risk of developing PD, but does not lead to GD.

In conclusion, this study supports the hypothesis that the mechanisms associated with individual *GBA* mutations may be multiple and provides evidence that the E326K mutation does not behave in the same way as common loss of function mutations, L444P and N370S.

## Materials and Methods

### Cell lines

The majority of human fibroblast cell lines were taken from the Schapira laboratory cell line bank. Participants gave informed consent and the collection of skin biopsies was approved by the Royal Free Research Ethics Committee (REC number 10/H0720/21) and the Great Ormond Street Hospital Ethics Committee. Cell lines GM10905 (Coriell Cat# GM20272, RRID:CVCL_0R42) and GM20272 (Coriell Cat# GM20272, RRID:CVCL_0R42) (L444P/L444P) were purchased from Coriell Institute cell repository. ND41016 (Subject ID: NDS00203) (NHCDR Cat# ND41016, RRID:CVCL_EZ71) (E326K/E326K) was purchased from NINDS Stem Cell Catalogue. Cell lines UCL-CTRL001 (WT/WT) (RRID:CVCL_B7T4); UCL-CTRL002 (WT/WT) (RRID:CVCL_B7T5); UCL-CTRL003 (WT/WT) (RRID:CVCL_B7T6); UCL-YCTRL001 (WT/WT) (RRID:CVCL_B7T8); UCL-E001K (WT/E326K) (RRID:CVCL_B7T9); UCL-N001S (N370S/N370S) (RRID:CVCL_B7TA) and UCL-N002S (N370S/N370S) (RRID:CVCL_B7TB) were generated in the Schapira laboratory and are being deposited to ATCC and will be available by final publication. Fibroblast cell line 7301 was obtained from the MRC Centre for Neuromuscular Diseases Biobank London, supported by the National Institute for Health Research Biomedical Research Centres at Great Ormond Street Hospital for Children NHS Foundation Trust and at University College London Hospitals NHS Foundation Trust and University College London. Details of the fibroblast cell lines used in this study can be found in Supplemental Table 1. Parental SH-SY5Y cells were purchased from ATCC (ATCC Cat# CRL-2266, RRID:CVCL_0019).

### Fibroblast cell culture

Fibroblasts were cultured in Dulbecco’s modified eagle media (DMEM) 4500 (mg/L) growth medium supplemented with Glutamax (Invitrogen), 10% foetal bovine serum (FBS), non-essential amino acids (NEAA: 0.1 mM of: glycine, L-alanine, L-asparagine, L-aspartic acid, L-glutamic acid, L-proline and L-serine) and penicillin/streptomycin antibiotic cocktail (50ng/ml) at 37°C and 5% CO2 (dx.doi.org/10.17504/protocols.io.rm7vzy542lx1/v1). Analyses were carried out at low passages, and disease and control cultures were matched for passage number.

### Generation of SH-SY5Y cell lines

Proliferating SH-SY5Y neuroblastoma cell lines were cultured as previously described (33) (dx.doi.org/10.17504/protocols.io.bp2l617jzvqe/v1). SH-SY5Y cells were transfected with a pcDNA3.1 vector plasmid containing wild-type, E326K, L444P or N370S *GBA* cDNA. Mutations were introduced by site directed mutagenesis, according to the manufacturer’s guidelines (Agilent Technologies QuikChange II Site-Directed Mutagenesis Kit) (https://www.agilent.com/cs/library/usermanuals/public/200523.pdf). Stable transfection was performed using the XtremeGENE reagent for 72 hours, and 400 µg/ml G418 antibiotics used as the selection marker (dx.doi.org/10.17504/protocols.io.rm7vzy54rlx1/v1). Colonies were selected and expanded for routine culture in growth media supplemented with G418. For analysis two clones per genotype were used.

### SDS-PAGE and Western blotting

Cells were lysed in 1% (v/v) Triton X-100 in phosphate buffered saline (PBS) lysis buffer with protease and phosphatase inhibitors. Cell lysates containing 10 – 30 µg of protein were electrophoresed with NuPage™ 4-12% Bis-Tris protein gels. Proteins were transferred to a PVDF membrane, blocked in 10% milk and treated with primary and secondary antibodies in 5% milk. For analysis of alpha-synuclein, an additional step was added prior to blocking to fix the membrane with 4% PFA (w/v) and 0.01% (v/v) glutaraldehyde for 30 minutes. Antibody binding was detected using the GE Healthcare Amersham™ electro-chemi-luminescence (ECL)™ Prime Western Blotting Detection Reagent (dx.doi.org/10.17504/protocols.io.261genwyyg47/v1). The following antibodies were used: GBA clone 2e2 (AP1140 Millipore, dilution 1:1000); GRP78/BiP (ab21685 Abcam, dilution 1:1000); monomeric alpha-synuclein (MJFR1 ab138501 Abcam, dilution 1:1000); LAMP1 (NB120-19294 Novus, dilution 1:1000); β-actin (ab8227 Abcam, 1:10000).

### Triton X-100 extraction of soluble and insoluble alpha-synuclein

SH-SY5Y cells were trypsinised and lysed in 1% (v/v) Triton X-100, 50 mM Tris, pH 7.5, 750 mM NaCl, 5 mM EDTA, 4 units RQ1 RNase-free DNase (Promega), protease and phosphatase inhibitor mix (Halt) on ice for 15 minutes. The lysate was passed through a 23G needle and pelleted at 17,000 x *g* for 30 minutes at 4°C. Triton X 100 soluble fraction was collected and protein concentration measured using the BCA protein assay. Insoluble pellets were resuspended in 2% (w/v) SDS, 8 M urea, 10 mM Tris, pH 7.5, 4 units RQ1 RNase-free DNase, protease and phosphatase inhibitor mix and incubated for 15 minutes at room temperature. Debris was removed by centrifugation at 17,000 x *g* for 30 minutes at 4°C (dx.doi.org/10.17504/protocols.io.6qpvr6p1ovmk/v1).

### Quantitative real-time PCR of mRNA

RNA was isolated from cells using the QIAGEN RNeasy mini kit (https://dx.doi.org/10.17504/protocols.io.ssbeean). Quantitative analysis of mRNA was performed as previous (99) (dx.doi.org/10.17504/protocols.io.q26g74xoqgwz/v1). GAPDH was amplified as the reference mRNA. Relative gene expression was calculated using the ΔCT method. All results obtained were from the evaluation of two technical duplicates of three independent experiments.

### Total cellular lysosomal enzyme assays

Cells were lysed in 1% (v/v) Triton X-100 in PBS with protease and phosphatase inhibitors. Activity was measured as described previously (10) GCase activity was measured in McIIvaine buffer at pH 5.4 in the presence of 22 mM sodium taurocholate (NaT) and at pH 4.5, with 5 mM methylumbelliferyl-β-D-glucopyranoside (M-Glu) substrate (dx.doi.org/10.17504/protocols.io.n2bvj625nlk5/v1). β-Galactosidase and β-Hexosaminidase activity was measured in McIIvaine buffer at pH 4.1 with 1 mM 4-Methylumbelliferyl β-D-galactopyranoside and 2 mM 4-Methylumbelliferyl N-acetyl-β-D-glucosaminide substrates, respectively (dx.doi.org/10.17504/protocols.io.kqdg3p8r7l25/v1).

### Real-time lysosomal GCase assay

Lysosomal GCase activity, but not ER and Golgi-resident GCase activity, can be measured in live cells by using 5-(pentafluorobenzoylamino) fluorescein Di-β-D-glucopyranoside (PFB-FDGlu; ThermoFisher) as a substrate that is taken up into acidic vesicles only, where it fluoresces upon catalysis with GCase (46) (dx.doi.org/10.17504/protocols.io.eq2lyn8mqvx9/v1). . Fibroblasts were grown to 70-90% confluency and the real-time GCase activity assay was performed as described previously (99). Conduritol-B-epoxide (CBE)-sensitive initial rate was calculated and normalised to protein concentration in the well. Cells were measured in triplicates.

### Endo H analysis

Fibroblasts were lysed in 1% (v/v) Triton X-100 in PBS and protein concentration determined by a BCA assay. Digestions were performed according to the manufacturer’s instructions (New England BioLabs #P0702L and #P0704S) (dx.doi.org/10.17504/protocols.io.8epv59ew4g1b/v1 and (dx.doi.org/10.17504/protocols.io.cqfvtm). For wild-type, E326K and N370S mutant lines, 20 μg of total protein was denatured and for L444P mutants 60 μg of total protein was denatured. Following this, denatured protein sample was incubated for 2 hours at 37°C, with 500,000 units/ml of either Endo H or PNGase F and analysed by western blotting.

### Staining for lipid droplets

Cells were grown on coverslips until 50-80% confluent. Analysis was performed on SH-SY5Y cells cultured in normal media, as well as fibroblasts and SH-SY5Y cells treated for 5 hours with 100 µM oleic acid (OA) following over-night starvation in Opti-MEM (ThermoFisher). OA is a potent inducer of lipid droplet formation (55). Cells were treated with 250 µL BODIPY™ 493/503 (4,4-Difluoro-1,3,5,7,8-Pentamethyl-4-Bora-3a,4a-Diaza-*s*-Indacene) diluted in PBS at a final concentration of 1 mg/ml for 15 minutes in the dark at room temperature. Following incubation, coverslips were washed three times in PBS and mounted onto glass slides with 1 μg/mL DAPI CitiFluor for analysis on a Zeiss Axioplan microscope (QImaging wLS). Images were captured using Micro-Manager software and analysed using ImageJ software with a macro to calculate the number of lipid droplets and cell area (dx.doi.org/10.17504/protocols.io.q26g74xqqgwz/v1).

### Statistics

Graphs were made using GraphPad Prism version 9.3.1 and data expressed as MEAN ± standard error of the mean (SEM). Statistical significance was determined by one-way ANOVA and Tukey’s post hoc or two-way ANOVA and Tukey’s post hoc test using GraphPad Prism version 9.3.1. Data distribution was assumed as normal without formal testing.

## Acknowledgements

The MRC Centre for Neuromuscular Diseases Biobank London, supported by the National Institute for Health Research Biomedical Research Centres at Great Ormond Street Hospital for Children NHS Foundation Trust and at University College London Hospitals NHS Foundation Trust and University College London, is acknowledged for providing the fibroblast cell line 7301.

## Abbreviations

PD: Parkinson disease
GCase: Glucocerebrosidase
ER: Endoplasmic reticulum
GBA-PD: GBA mutation-associated Parkinson disease
GlcCer: Glucosylceramide
GD: Glucosylsphingosine – GlcSph Gaucher disease
ALP: Autophagic-lysosomal pathway
UPR: Unfolded protein response
ROS: Reactive oxygen species
DMEM: Dulbecco’s modified eagle media
FBS: Foetal bovine serum
NEAA: Non-essential amino acids
PBS: Phosphate buffered saline
ECL: Electro-chemi-luminescence
CBE: Conduritol-B-epoxide
OA: Oleic acid
SEM: standard error of the mean

