## Supplemental Figure 1 for "The *GBA* variant E326K is associated with alpha-synuclein aggregation and lipid droplet accumulation in human cell lines"

(A)

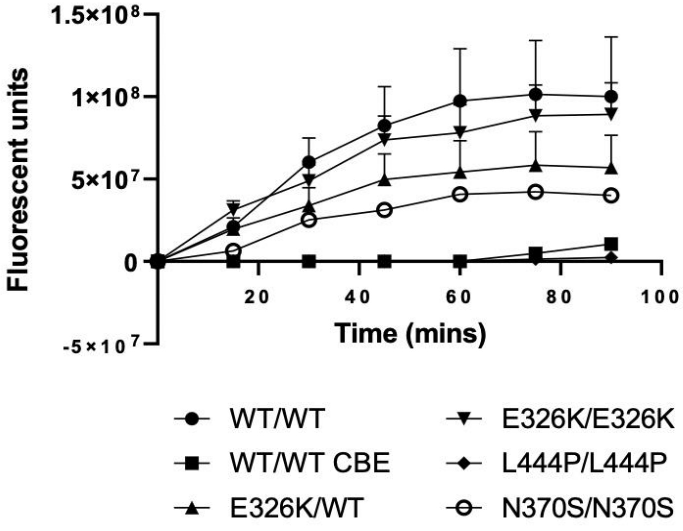


(B)

| Genotype | Linear trendline equation (y=mx) |
|---|---|
| WT/WT | y=18403371x |
| WT/E326K | y=11332758x |
| E326K/E326K | y=16823338x |
| L444P/L444P | y=0x |
| N370S/N370S | y=7128687x |
| WT/WT + CBE | y=0x |

**Supplementary Figure 1. Activity of lysosomal GCase in patient fibroblast lines. (A)** Fibroblast cell lines were incubated with PFB-FDGluc substrate for 1 hour at 37°C. Following washing, fluorescence was measured every 15 minutes for 90 minutes. **(B)** The initial linear rate of each reaction was calculated between time 0 and time 45 minutes and initial rate equations displayed in the table. Raw data can be found at: 10.5281/zenodo.6553597
