## Supplemental Table 1 for "The *GBA* variant E326K is associated with alpha-synuclein aggregation and lipid droplet accumulation in human cell lines"

**Supplementary Table 1. Characteristics of fibroblast cell lines.** For fibroblast analysis cell lines used were: UCL-CTRL001, UCL-CTRL002, UCL-CTRL003, 7301, UCL-YCTRL001, UCL-E001K, ND41016, GM10905, GM20272, UCL-N001S and UCL-N002S.

| Cell line | GBA genotype | Phenotype | Gender | Age |
|---|---|---|---|---|
| UCL-CTRL001 | WT/WT | Unaffected | F | 58 |
| UCL-CTRL002 | WT/WT | Unaffected | M | 53 |
| UCL-CTRL003 | WT/WT | Unaffected | F | 73 |
| 7301 | WT/WT | Unaffected | F | 14 |
| UCL-YCTRL001 | WT/WT | Unaffected | M | 1 |
| UCL-E001K | WT/E326K | PD | M | 50 |
| ND41016 | E326K/E326K | PD | M | 52 |
| GM10905 | L444P/L444P | Type I GD | M | 7 |
| GM20272 | L444P/L444P | Type II GD | M | Child |
| UCL-N001S | N370S/N370S | GD | F | 58 |
| UCL-N002S | N370S/N370S | GD | M | 76 |
